# Determining the targeting specificity of the selective peroxisomal targeting factor Pex9

**DOI:** 10.1101/2022.02.02.478779

**Authors:** Eden Yifrach, Markus Rudowitz, Luis Daniel Cruz-Zaragoza, Asa Tirosh, Zohar Gazi, Yoav Peleg, Markus Kunze, Miriam Eisenstein, Wolfgang Schliebs, Maya Schuldiner, Ralf Erdmann, Einat Zalckvar

## Abstract

Targeting proteins to their correct cellular location is a fundamental process that allows them to carry out their cellular functions. Peroxisomes utilize two paralog targeting factors, Pex5 and Pex9, for proteins with a Peroxisomal Targeting Signal 1 (PTS1). However, in spite of their similarity, Pex9 targets only a subset of Pex5 cargo proteins. Here, we studied the properties that facilitate the targeting specificity of Pex9, both by unbiased screens and by site-directed mutagenesis of the PTS1 motifs of either binders or non-binders. We find that the binding specificity of Pex9 is largely determined by the hydrophobic nature of the amino acid preceding the PTS1 tripeptide of its cargos. This is in line with structural modeling of the PTS1-binding cavity of the two factors, showing that while Pex5 has large negative electrostatic patches at the area surrounding the PTS1 binding cavity, Pex9 is mostly hydrophobic. Our work outlines the mechanism by which targeting specificity is achieved, enabling dynamic rewiring of the peroxisomal proteome in changing metabolic needs.

## Introduction

Targeting proteins to their correct cellular location is a fundamental process for the life of every organism. In eukaryotes, nearly all proteins are synthesized by ribosomes in the cytosol and must be targeted to their designated compartment to function properly (Hegde and Zavodszky, 2019). Targeting to the appropriate compartment allows a protein to form the necessary interactions with its partners and participate in biological networks such as signaling and metabolic pathways (Laurila and Vihinen, 2009). The efficacy/accuracy of the targeting machinery also prevents the protein from mislocalizing to other organelles or aggregating in the cytosol. For these reasons, mutations affecting the targeting of individual proteins or the targeting machinery itself can have severe functional consequences on cells and cause disease (Schaeffer et al., 2014).

While cellular compartments have distinct molecular machineries for directed transport, the shared feature of these processes involves the recognition of a targeting signal within the nascent protein by a destination-specific targeting factor. Thus, the fidelity of cellular spatial organization relies critically on the specificity and efficiency by which signals on the targeted proteins are recognized by their associated targeting factors (Hegde and Zavodszky, 2019; Aviram and Schuldiner, 2017).

One of the organelles that has a complex and fascinating targeting machinery is the peroxisome. Peroxisomes perform and regulate a myriad of metabolic activities, such as degradation of fatty acids and regulation of redox homeostasis (Islinger et al., 2018), for which the proper import of lumenal enzymes is essential. To import peroxisomal matrix (lumen) proteins, two constitutive targeting factors, Pex5 and Pex7, recognize proteins with a Peroxisomal Targeting Signal (PTS) type I and type II, respectively, and shuttle them to the organelle (Walter and Erdmann, 2019). We previously identified an additional targeting factor, Pex9, which is expressed in yeast under specific metabolic conditions and targets only a subset of PTS1 proteins (Yifrach et al., 2016; Effelsberg et al., 2016).

The PTS1 is defined as a tripeptide at the C terminus (C’) of the protein with additional amino acids upstream also contributing to the binding (Lametschwandtner et al., 1998; Stanley et al., 2006; Fodor et al., 2012; Hagen et al., 2015; DeLoache et al., 2016; Hochreiter et al., 2020). The ultimate yeast PTS1 tripeptide contains a small uncharged residue (serine (S)/ alanine (A)), then a positively charged residue (arginine (R)/ lysine (K)/ histidine (H)), and at the extreme C’ a leucine (L) or phenylalanine (F) (Brocard and Hartig, 2006). We found that Pex9 targets the three enzymes Mls1, Mls2, and Gto1, all containing a classical PTS1 tripeptide (Yifrach et al., 2016; Effelsberg et al., 2016). Pex9 presumably prioritizes these specific enzymes that are required in peroxisomes under fatty-acid-dependent growth thus enabling dynamic rewiring of peroxisomes in response to metabolic needs. But how does Pex9, which is a paralog of Pex5, recognize only a subset of PTS1 proteins, and what defines the targeting specificity of Pex9?

We took four approaches to better understand the targeting specificity of Pex9. First, we looked for additional Pex9 cargos amongst the recently identified yeast peroxisomal proteins (Yifrach et al., 2021) and used one such new protein alongside the known cargo to align the PTS1 and look for similar patterns. Second, we performed site-directed mutagenesis on the PTS1 sequences of known Pex9 and Pex5 cargos and looked at their effect on Pex9 dependent targeting to peroxisomes and physical binding. Third, we modeled the PTS1-binding domain of Pex9 and Pex5 to distinguish features on the area surrounding the binding cavity or the cavity itself of each cargo factor that give preference and specificity to some PTS1 sequences. And finally, we performed an unbiased screen with variable PTS1 sequences to validate our findings of the properties that enable Pex9 recognition.

Using the different approaches, we found that Pex9 prefers hydrophobic and negatively charged residues upstream to the PTS1 tripeptide. This is in contrast to Pex5 which was previously shown to prefer positively charged residues in these positions (DeLoache et al., 2016). This was supported by our modeling of the surface of the PTS1 binding domain that showed that the area surrounding the PTS1 binding cavity of Pex9 is mostly hydrophobic and with positively charged edges compared to the more negative surface of Pex5. These distinct features are another example of how complex and intricate the targeting landscape of peroxisomes is to allow differential targeting, which enables the peroxisome to rewire its function according to metabolic needs.

## Results

### Expanding the cargo range of Pex9 using a microscopy screen

To align the PTS1 of Pex9 cargo to find similar patterns it is important to have the biggest possible sample size. However, to date, only three Pex9 cargos have been found. Therefore, we sought to find additional proteins that can be targeted by Pex9. To this end, we performed a microscopic screen on a collection of ~40 yeast strains representing a set of recently identified yeast peroxisomal proteins (Yifrach et al., 2021), which were never tested for targeting by Pex9 (Fig. 1A). All proteins in this collection harbor a Green Fluorescent Protein (GFP) tag at the amino terminus (N’) for visualization and to allow their C’ to be exposed in case they have a PTS1. To examine the dependence of the peroxisomal proteins on Pex9, we used an automated mating procedure to insert several genetic traits to each strain: I) a peroxisomal marker, Pex3-mCherry, II) a deletion of the main PTS1 targeting factor (*Δpex5*) and III) a constitutive expression of *PEX9* to enable visualization of its function in glucose-containing media, as was done previously (Yifrach et al., 2016). We imaged the entire collection using a high content screening setup and looked for proteins that co-localize with peroxisomes when only Pex9 is expressed as a PTS1 targeting factor. Following analysis of all strains, we were able to identify two proteins that co-localized with peroxisomes in this condition – Fsh3, a newly-identified peroxisomal lipase (Yifrach et al., 2021), and Afr1, a protein required for the formation of pheromone-induced projection in yeast (Konopka, 1993) (Fig. 1B).

**Figure 1.**
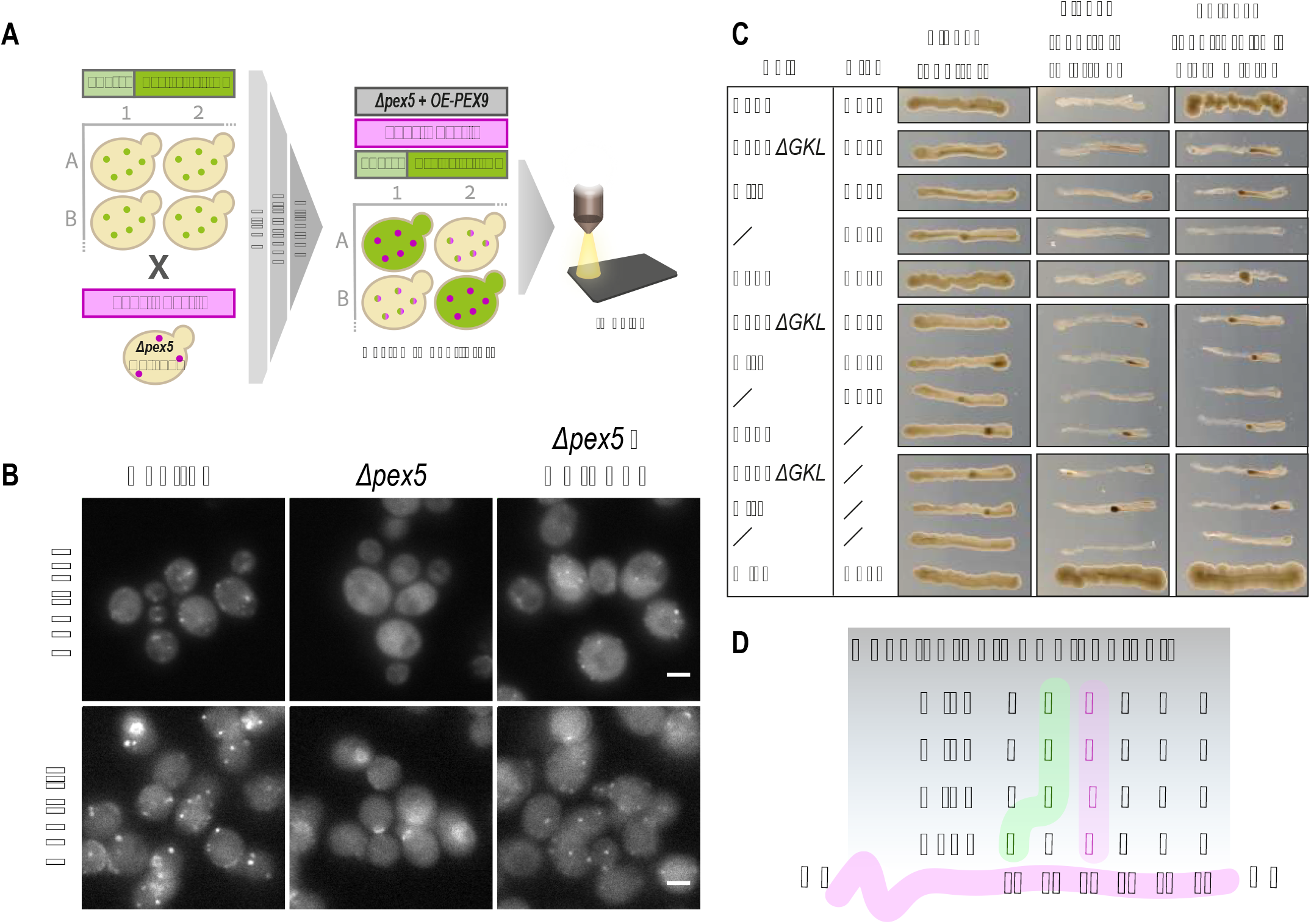
Expanding the cargo range of Pex9 using a microscopy screen. (A) A query strain constitutively expressing Pex9 as the sole PTS1 targeting factor *(Δpex5 + TDH3pr-PEX9)* and a peroxisomal marker (Pex3-mCherry) was genomically integrated into a yeast N′ GFP collection of recently identified peroxisomal proteins (Yifrach et al., 2021) by utilizing an automated mating procedure. Then, fluorescence microscopy was applied to identify proteins that co-localize with the peroxisomal marker when the cells grew on media containing either glucose or oleate as the carbon source. (B) Fsh3 and Afr1 both co-localize with peroxisomes when Pex9 is constitutively expressed and *PEX5* is deleted suggesting they are newly-identified cargos of Pex9. (C) Yeast 2-hybrid (Y2H) assays demonstrate that Fsh3 interacts with Pex9 in a PTS1 dependent manner, but Afr1 does not. (D) All Pex9 cargos show similar amino acid properties at positions −4 (hydrophobic residue L or I) and −5 (negatively charged residue D) from the C’ of the protein.

To assess their capacity to bind Pex9, we used a Yeast-2-hybrid (Y2H) assay. We found that Fsh3, but not Afr1, physically interacts with Pex9 (Fig. 1C). Afr1 does not seem to contain a PTS1 at its C’ (the last six amino acids of the protein are FTHYLI), which altogether suggests that Afr1 could piggyback on another Pex9 cargo that contains a PTS1 similarly to a previously shown Pex5 cargo (Gabay-Maskit et al., 2020). In addition, we show that the binding of Fsh3 to Pex9 depends on the PTS1 tripeptide of Fsh3 and that neither Fsh3 nor Afr1 interact in Y2H assays with Pex5. Put together, this suggests that Fsh3 is an additional direct cargo for Pex9.

After we expanded our cargo list, we could better assess unique sequence features. We compared the context of the PTS1 (the amino acids upstream to the PTS1 tripeptide) of the four known Pex9 cargos Mls1, Mls2, Gto1, and Fsh3 (Fig. 1D). We noticed that all cargos have a hydrophobic residue one amino acid upstream to the tripeptide (position −4 from the C’) and a negatively charged residue in position −5 or −6 from the C’. This similarity suggests that these features are important for the recognition by Pex9.

### Mutagenesis on the PTS1 context of known cargos affects Pex9 targeting

To explore whether the negatively charged residue in the PTS1 context of the Pex9 cargos plays an important role in their recognition, we substituted the charged residues at this position in various cargo proteins and looked for changes in the targeting ability. First, we constructed an integration plasmid containing a GFP fused at its C’ to the last 10 amino acids of Fsh3. We transformed this construct into the genome of a strain that constitutively expresses only Pex9, but not Pex5 (*Δpex5* + *TDH3pr-PEX9*). We validated that the PTS1 motif of Fsh3 is sufficient to target the GFP to peroxisomes by Pex9 (Fig. 2A, wtPTS1). Then, we mutated the negatively charged residue aspartate (D) in position −6 of the PTS1 motif to A that has no charge. We indeed observed a reduction in the peroxisomal localization of the protein (Fig. 2A and Fig. 2B, D(−6)A) but not a complete loss of the peroxisomal localization.

**Figure 2.**
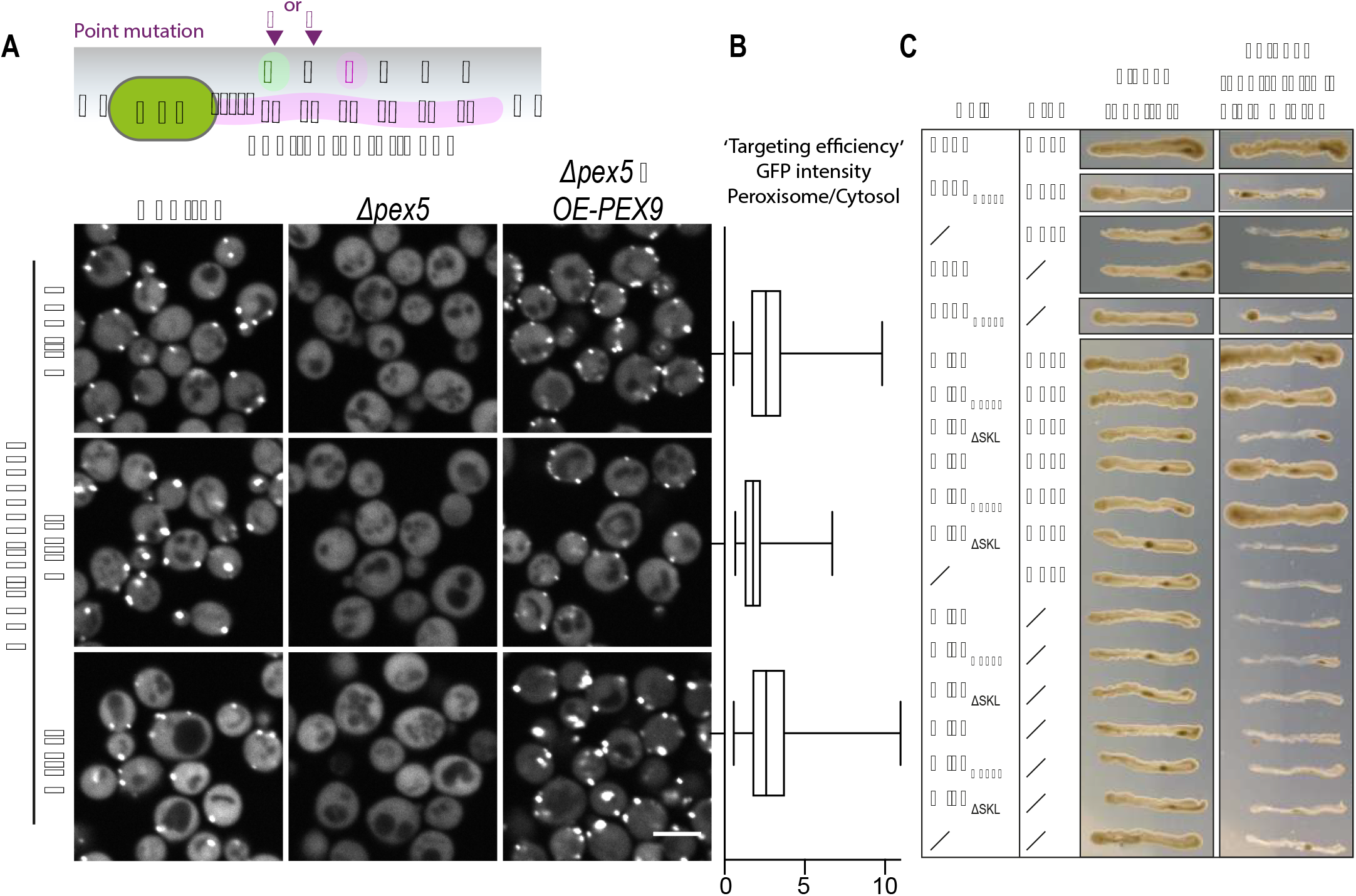
Mutagenesis on the PTS1 context of known cargos affects Pex9 targeting. (A) Directed point mutagenesis was applied on an integration plasmid containing a GFP fused at its C’ to the last 10 amino acids of Fsh3 to substitute position −6 or −5 from aspartic acid (D) or serine (S) to alanine (A). While the substitution of D to A at position −6 showed reduced GFP localization to peroxisomes, the S to A substitution at position −5 showed enhanced GFP localization in peroxisomes when the cells express Pex9 as the sole PTS1 targeting factor. (B) Image analysis quantification of the peroxisomal signal relative to cytosolic one (indicative of ‘targeting efficiency’) for the different constructs when Pex9 is expressed as the sole PTS1 targeting factor. (C) Y2H assays show that a D to lysine (K) substitution in the PTS1 context of Fsh3 obliterate the interaction with Pex9, but a similar substitution of D to K in Mls1 and Mls2 does not affect the interaction with Pex9, suggesting that D in the context of the PTS1 is not the sole nor definitive determinant for Pex9 cargo recognition.

Furthermore, when we performed Y2H assays and mutated the D to a positively charged residue, K, it obliterated the interaction of Fsh3 with Pex9 (Fig. 2C). However, when we mutated the conserved D to K in the Pex9 cargos Mls1 and Mls2, the proteins could still interact with Pex9. These data suggest that the negatively charged residue in position −6 or −5 of the PTS1 plays a role in some proteins but is not the sole nor definitive determinant for Pex9 cargo recognition.

Since factor-specific targeting requires that a cargo binds Pex9 in an enhanced affinity and that it binds Pex5 with reduced affinity, we also assayed the effect of the above-mentioned mutations on Pex5 binding. Interestingly, while the wild-type Mls1 and Mls2 (harboring a D at position −5) did not interact with Pex5, the D to K exchange in Mls1 and Mls2 promoted the interaction with Pex5 (Fig. S1). These observations are in agreement with previous data showing that positively charged residues in the PTS1 context enhance targeting by Pex5 (DeLoache et al., 2016). We speculate that the D in position −5 or −6 of the Pex9 cargos serves to reduce capture by Pex5, rather than to enhance binding of Pex9.

Next, we tested whether a mutation closer to the PTS1 tripeptide affects the targeting ability of Pex9. We changed S to A in position −5 of Fsh3 and checked how well the GFP-PTS1 construct localizes to peroxisomes. Although the basal targeting of the native PTS1 construct was already quite good, we were able to observe enhanced peroxisome to cytosol signal (which we define as targeting) in the mutated construct compared to the native PTS1 (Fig. 2A and Fig. 2B, S(−5)A). S is a polar amino acid, while A is non-polar. This suggests that Pex9 prefers a non-polar amino acid in this position.

### Exploring Pex9 targeting specificity using natural non-binders

Intrigued by the ability to enhance import by Pex9, we decided to mutate Pex9 non-binder PTS1 motifs to potentially induce import, which will facilitate visualization as there will be no background targeting. To start with a Pex9 non-binding PTS1 we first tested the motifs of several PTS1 proteins that are not localized to peroxisomes when Pex9 is expressed and Pex5 is absent. We tested constructs containing a GFP fused at its C’ to the last ten amino acids of Mdh3, Cat2, or Lys1. To our surprise, we found that the last 10 amino acids of both Mdh3 and Cat2 were sufficient to co-localize the GFP to peroxisomes in a Pex9 dependent manner (Fig. 3A and Fig. 3B), although the full-length proteins are not Pex9 cargos. We used a Y2H assay to validate the microscopy-based observation and showed that the full-length protein Mdh3 does not interact with Pex9, while a peptide consisting of only the last 10 amino acids of Mdh3 does (Fig. 3C). In line with this, we found that the Pex9 cargos, Fsh3, and Mls1 full-length proteins, do not interact with Pex5 in Y2H, but peptides consisting of only their last 10 amino acids can bind Pex5 (Fig. 3C). This striking observation suggests that additional parameters in the full protein prevent, or reduce, the interaction with the inappropriate factor and block targeting even when the PTS1 by itself could enable binding.

**Figure 3.**
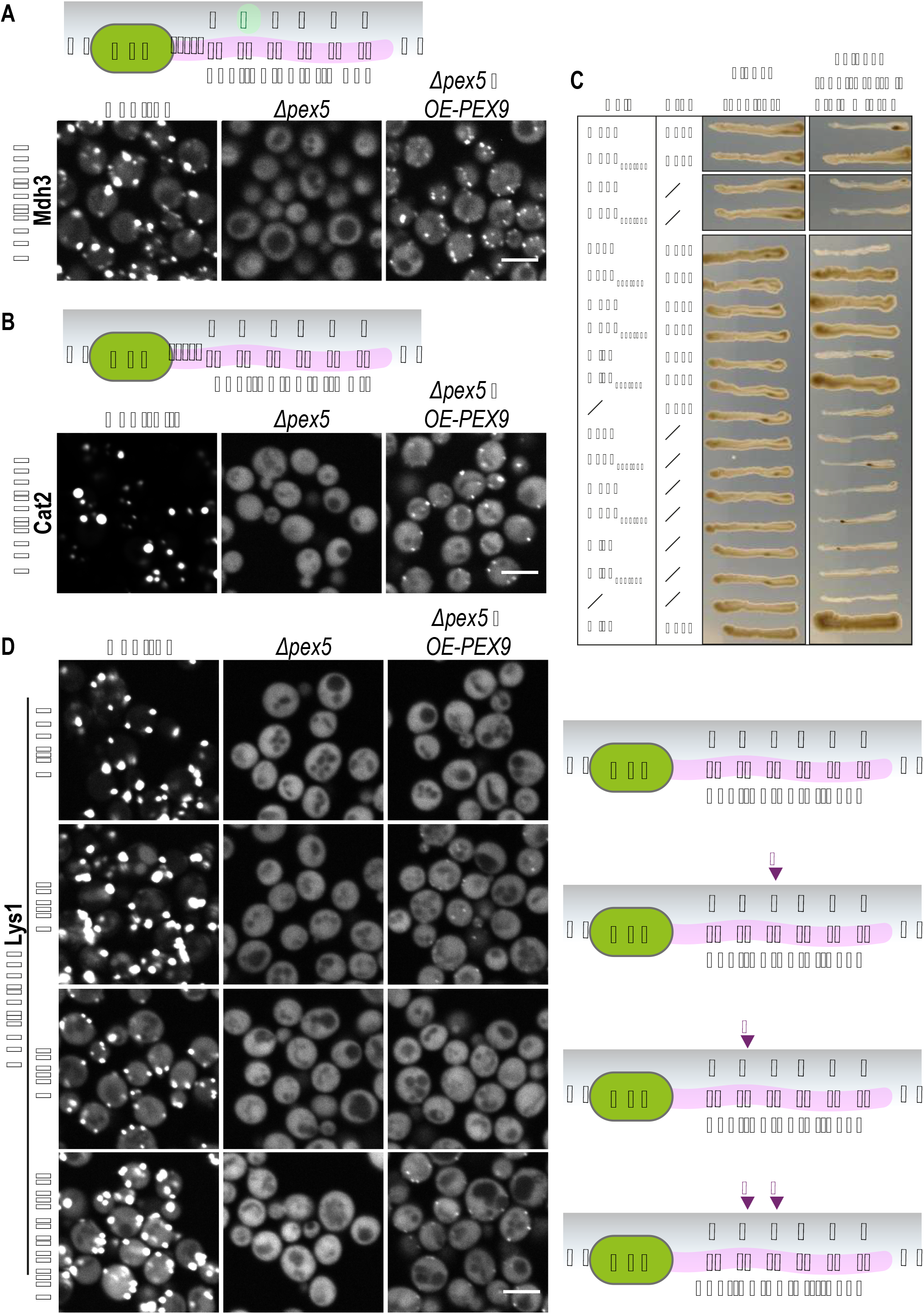
Exploring Pex9 targeting specificity using natural non-binders. Integration plasmids containing a GFP fused at its C’ to the last 10 amino acids of (A) Mdh3 or (B) Cat2 show that the last 10 amino acids of both Mdh3 and Cat2 were sufficient to co-localize the GFP to peroxisomes in a Pex9 dependent manner, although the full-length proteins are not cargos of Pex9. The asterisk indicates that GFP-last 10aaCat2 Control (in B) had a higher signal compared to the other images in the panel, hence a different image contrast was used. (C) Y2H assays validate that the last 10 amino acids of Mdh3 (aa334-343) can interact with Pex9 despite the fact that the full-length protein does not. Similarly, the last 10 amino acids of the Pex9 cargos Fsh3 (aa257-266) and Mls1 (aa545-554) interact with Pex5, more strongly than the full-length proteins. This suggests that additional parameters in the full protein prevent, or reduce, the interaction with the inappropriate targeting factor. (D) An integration plasmid containing a GFP fused at its C’ to the last 10 amino acids of Lys1 shows that the native PTS1 of Lys1 was not sufficient to co-localize the GFP to peroxisomes in a Pex9 dependent manner. Directed mutagenesis substitution of positions −5 and −4 to a negatively charged residue D and/or a non-polar residue leucine (L), show that the L mutation enabled a weak GFP peroxisomal localization, demonstrating that a single non-polar residue at position −4 is sufficient to mediate Pex9 specificity.

We continued to work on the last 10 amino acids of Lys1 that were not sufficient to mediate Pex9 dependent targeting (Fig. 3D, upper panel). To enhance Pex9 binding, we replaced the amino acids at positions −5 and −4 with a negatively charged residue D and a non-polar residue L, either alone or in combination. While the D mutation had no visible effect on the localization of the GFP, the constructs with the L mutation showed a weak peroxisomal signal, demonstrating that a single amino acid change to a non-polar residue at position −4 is sufficient to allow binding to Pex9.

Overall, our data imply that the PTS1 targeting specificity of Pex9 is largely dependent on the presence of a non-polar, hydrophobic residue in the vicinity of the PTS1 tripeptide. Moreover, we suggest that the negatively charged residue at positions −5 or −6 of the PTS1 has two roles – to enhance the binding of specific cargo proteins to Pex9 and to prevent binding to Pex5.

### Pex9 molecular modeling support hydrophobicity in the PTS1 binding area as the main determinant of Pex9 binding

Our targeted approach highlighted the preference of Pex9 for a hydrophobic residue preceding the PTS1 tripeptide. This distinguishes Pex9 from Pex5, which is known to prefer positively charged residues in this context (DeLoache et al., 2016). To uncover the structural features that differentiate the two targeting factors in their binding preference, we constructed a model structure of the Pex9 PTS1-binding domain in a similar manner to the modeling of Pex5, as described previously (Gabay-Maskit et al., 2020) (Fig. 4A). The models highlight that the Pex5 surface around the PTS1 binding cavity has large negative electrostatic potential patches (red), while the surface of the cognate Pex9 regions appears more hydrophobic (white). This is especially obvious for the shallow cavity that binds the amino acid in position −2 and the bottom of the PTS1 binding cavity (green arrows, Fig. 4A). In addition, Pex9 has several positive electrostatic patches (blue) at the edge of the peptide-binding cavity.

**Figure 4.**
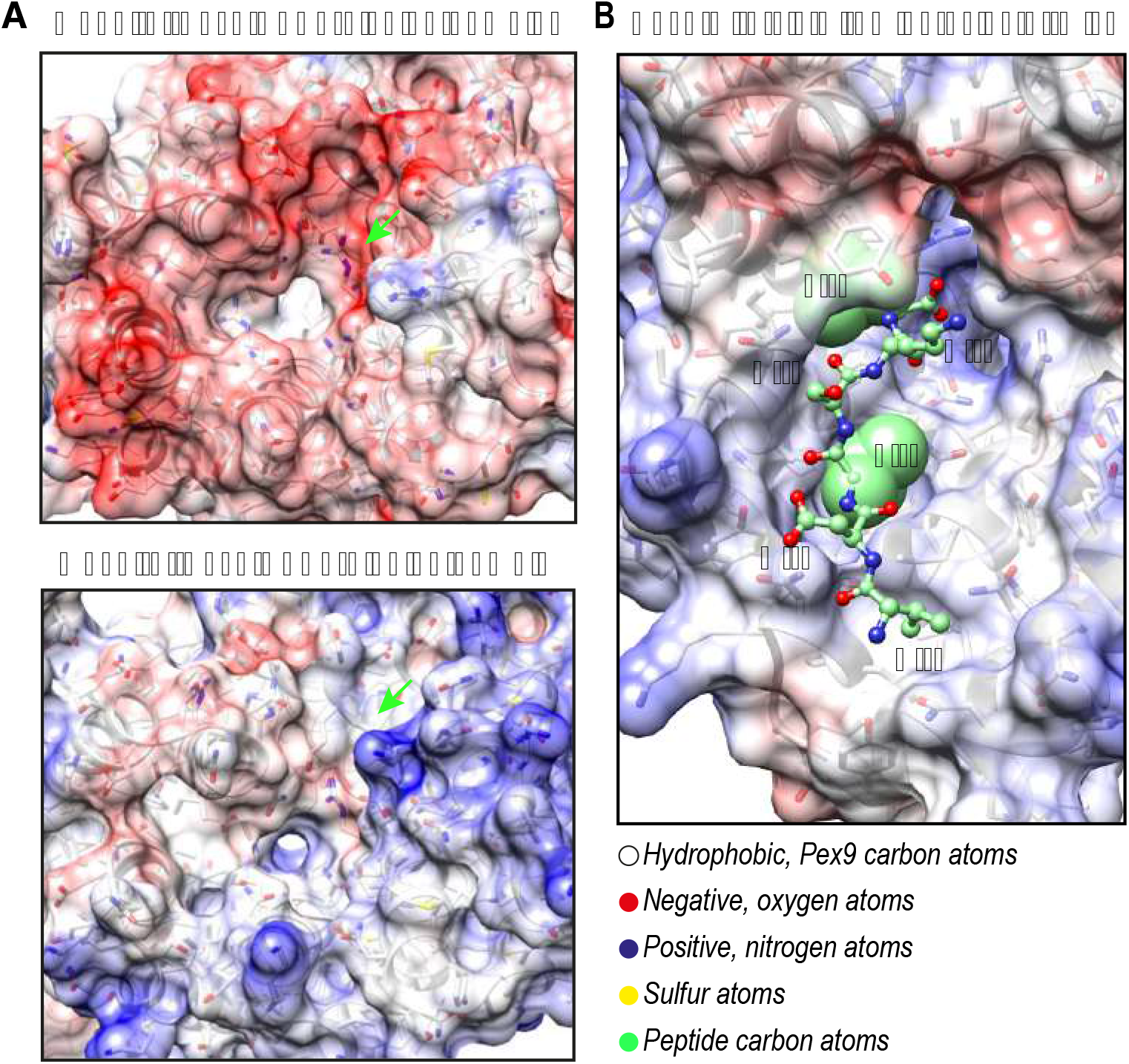
Pex9 molecular modeling support hydrophobicity in the PTS1 binding area as the main determinant of Pex9 binding. (A) Molecular modeling of the PTS1 binding domains of Pex9 and Pex5 indicate that the Pex5 surface around the PTS1 binding cavity has large negative electrostatic potential patches (red), while the surrounding area of the PTS1 cavity of the cognate Pex9 regions appears more hydrophobic (white) with several positive electrostatic patches (blue) at the edge of the peptide-binding cavity. Green arrows pointing at the shallow cavity that binds peptide residue −2. (B) Molecular dynamics simulations of complexes containing Pex9 with peptides consisting of six C’ amino acids of the known Pex9 cargos (Mls2 is presented) show that in less than 50ns, the peptide changed conformation and the sidechain of its −4 residue pointed toward the bottom of the PTS1 binding cavity and made tight contacts with hydrophobic Pex9 residues.

We then performed Molecular Dynamics (MD) simulations of a cargo factor (Pex5 or Pex9) with peptides consisting of six C’ amino acids of four PTS1 proteins that are Pex9 cargo: Mls1, Mls2, Gto1, and Fsh3. Previously we showed that the binding stability of a peptide to Pex5 could be estimated from the constancy of hydrogen bonds (H-bonds) between peptide backbone atoms and specific Pex5 residues, particularly H-bonds of peptide residues −1 and −3 (Yifrach et al., 2021). The Pex5 residues that form these H-bonds are conserved between Pex5 and Pex9 supporting the importance of the C’ tripeptide for binding. The context residues, however, behaved differently in the simulations for Pex5 and Pex9 complexes. In the starting structures of all the complexes, the side chain of peptide residue −4 pointed outwards, making either no, or little, contact with the cargo factor, as seen in the experimental structure that was used as the modeling template (Gatto et al., 2000). While in the Pex5 simulations this side chain remained mostly exposed, in the Pex9 simulations the peptide changed conformation in less than 50ns and the sidechain of its −4 residue pointed toward the bottom of the PTS1 binding cavity (Fig. 4B). This hydrophobic sidechain (L or isoleucine (I)) made tight contacts with hydrophobic Pex9 residues that replace more polar residues of Pex5 (e.g. tyrosine (Y)410, and A437 in Pex9 versus S507 and S534 in Pex5). Together with additional residues, they form a more hydrophobic surface at the entrance to the peptide-binding cavity of Pex9. These results strongly support the notion that Pex9 cargos are selected by the hydrophobic residue in position −4 of the PTS1.

### A systematic screen shows a correlation between Pex9-dependent peroxisomal localization and hydrophobicity of residue −4 of PTS1 sequences

To validate our findings in an unbiased manner and to investigate more variations of the PTS1 context, we took advantage of a library of integration plasmids containing randomized sequences preceding the ultimate PTS1 tripeptide S-K-L residues (DeLoache et al., 2016). This library was previously used to find the optimal Pex5 cargo sequence - the enhanced PTS1 import sequence. Each integration plasmid in this pooled library contains a Yellow Fluorescent Protein (YFP) linked at its C’ to a variable sequence of six amino acids preceding the SKL tripeptide. We transformed this YFP-Variable_sequence-SKL library genomically into a strain constitutively expressing Pex9 as the sole PTS1 targeting factor and picked single colonies into an arrayed format (Fig. 5A). Then, each strain was both imaged and the contained plasmid sequenced to match the ratio of its YFP-Variable_sequence-SKL peroxisome/cytosol localization to the identity of the variable amino acid sequence. We found a wide range of peroxisome/cytosol localization ratios, from very high to very low. Despite redundancy in sequences, we could retrieve 15 distinct linkers that correspond to the wide range of phenotypes (Fig. 5B). We manually validated two sequences from the screen by fusing them to the C’ of GFP (Fig. S2). We plotted the different PTS1 contexts according to their peroxisome/cytosol localization ratio and the hydrophobicity score (Trinquier and Sanejouand, 1998) of the residues at position −4 (Fig. 5C). This analysis emphasizes that the level of hydrophobicity at position −4 correlates with the level of peroxisomal targeting by Pex9 and validates our proposal that Pex9 cargos are selected by the hydrophobic residue in position −4 of the PTS1.

**Figure 5.**
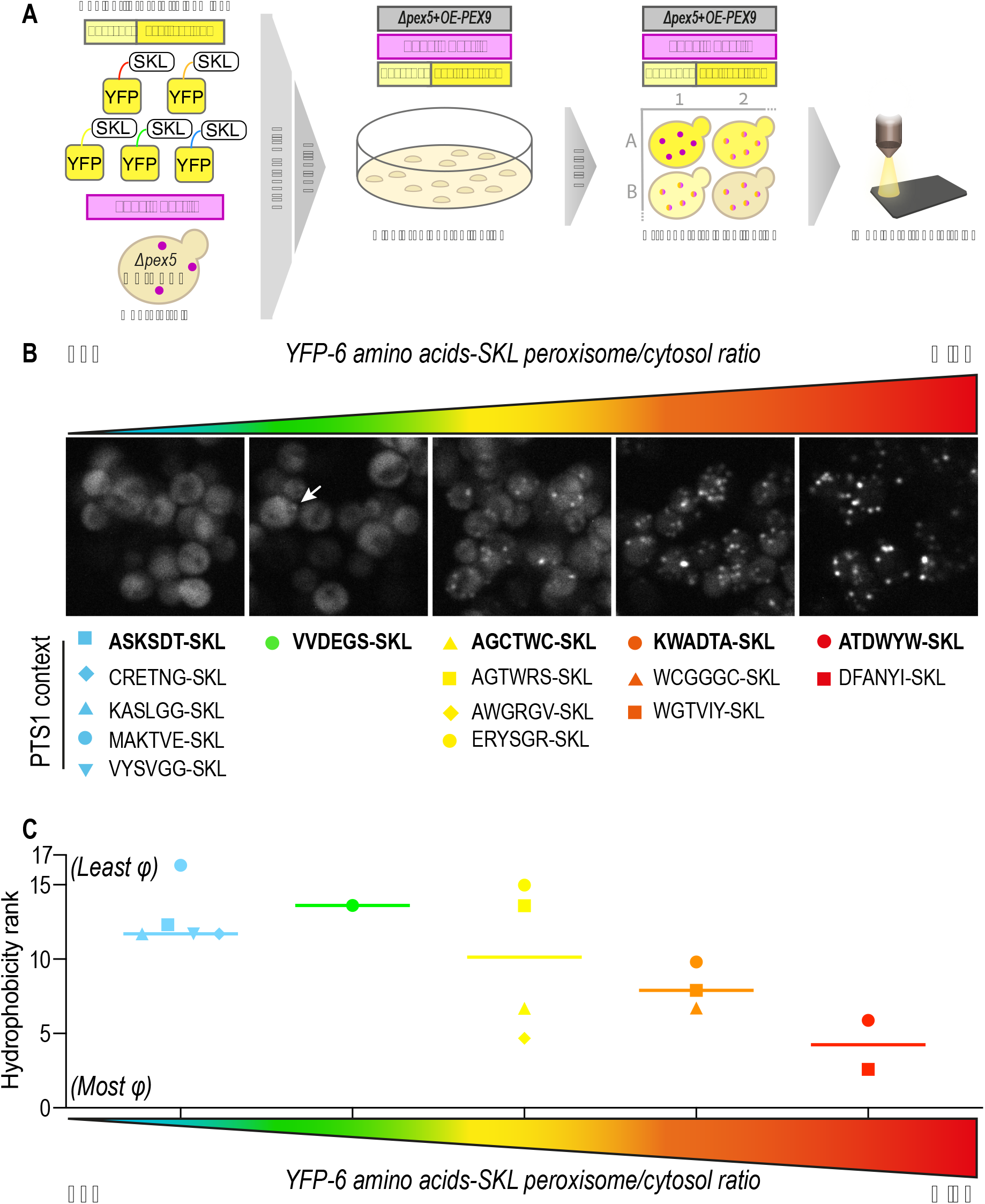
A systematic screen shows a correlation between Pex9-dependent peroxisomal localization and hydrophobicity of residue −4 of PTS1 sequences. (A) A pooled library of plasmids containing YFP fused to a stretch of six random amino acids followed by the tripeptide SKL was transformed into a strain constitutively expressing Pex9 as the sole PTS1 targeting factor *(Δpex5 + TDH3pr-PEX9)* and a peroxisomal marker (Pex3-mCherry). Cells from single colonies were imaged and the PTS1 context of each plasmid was sequenced. (B) Fifteen distinct PTS1 context sequences (the random stretches) were identified and matched to the peroxisome/cytosol localization ratio of their cognate YFP construct. The white arrow is pointing at a peroxisomal punctate. (C) Amino acid analysis shows that the YFP peroxisome/cytosol localization ratio correlates with the hydrophobicity score of the residue at position −4 from the C’ of each construct. This combined hydrophobicity rank (Trinquier and Sanejouand, 1998) gives low values for hydrophobic residues (most φ) and high values for hydrophilic residues (least φ). The analysis validates that Pex9 cargos are selected by the hydrophobic residue in position −4 of the PTS1.

## Discussion

‘Being in the right place at the right time’ is not simply a motto – when it comes to cellular proteins, it is an absolute necessity. In a cell, the right proteins must be shuttled from the cytosol to their destination compartment upon demand. When *S. cerevisiae* cells rely on fatty acid-containing media, peroxisomes become essential, as they are the sole organelles that break down fatty acids. Thus, peroxisomes must dynamically change their protein content according to the cell’s metabolic needs. To give targeting priority to essential enzymes, the metabolically regulated arm of the targeting machinery is activated. Both the targeting factor Pex9 and the Pex7 co-factor, Pex18, are upregulated to boost the targeting of specific PTS1 and PTS2 proteins, respectively (Effelsberg et al., 2015, 2016). Although Pex9 is a paralog of Pex5, it targets only a subset of Pex5 cargo proteins – Mls1, Mls2, and Gto1. Here we found two additional proteins that rely on Pex9, Fsh3, and Afr1.

How is the binding specificity achieved? Protein specificity is a function of both positive and negative selection, meaning the binding to one specific partner while *not* binding to others (Schreiber and Keating, 2011). Indeed, our work shows that some residues such as the −5/6 negative charge in context to the PTS1 tripeptide, can act either to enhance binding to Pex9 or weaken binding to Pex5. In this context, a recent study on the selectivity of the Golgi to Endoplasmic Reticulum (ER) retrieval signals shows that the KDEL receptors use a charge screening mechanism to differentiate between their cognate signals (Gerondopoulos et al., 2021). It was shown that the charge distribution across the surface of the KDEL receptor is used as an “antenna” for the initial signal capture and proofreading. We argue that an “antenna” like mechanism exists also for Pex5 and Pex9. Indeed, we observed that the electrostatic properties on the surfaces of the PTS1-binding cavities of Pex5 and Pex9 are significantly different. The surface of Pex5 has large negative electrostatic potential patches within and around the protein-binding cavity, while Pex9 in this region is mostly hydrophobic with several positive electrostatic patches at the edge of the binding cavity. We found that while the PTS1 motifs of non-binders can drive the targeting of GFP to peroxisomes, the full-length proteins are not interacting with the respective targeting factor, Pex5 or Pex9, in Y2H assays. This suggests that interaction interfaces outside of the PTS1 motif significantly contribute to the binding specificity. These binding interfaces can either attract the proteins to their cognate targeting factor and/or prevent their interaction with the inappropriate targeting factor at the initial stage of cargo proteins screening.

In addition, the properties of the PTS1 tripeptide itself and the preceding four or five amino acids, contribute to this positive and negative selection as well. We show by molecular modeling, a targeted experimental approach, and an unbiased screen that the amino acid at position −4 of the PTS1 has the most significant effect on binding selectivity by Pex9. The replacement of only a few residues from polar in Pex5 to hydrophobic in Pex9 forms a more hydrophobic region at the entrance to the peptide-binding cavity. Correspondingly, all Pex9 cargos contain a hydrophobic residue at position −4.

Our work exemplifies how changes in the binding cavity of two rather similar targeting factors lead to different cargo binding specificities. These findings expand the range of capabilities of how peroxisomes achieve exquisite targeting specificity – from the presence of multiple differentially regulated parallel pathways for targeting, through affinity-tuning of various PTS1s to provide priority targeting (Rosenthal et al., 2020), to post-translational modifications of Pex5 or its cargo proteins that alter its binding specificity. Each cargo has a unique targeting propensity that can also be regulated to enable the dynamic rewiring of protein content in peroxisomes upon demand and shows the beauty and complexity of the peroxisomal targeting machinery.

## Materials and methods

### Yeast strains and strain construction

All strains in this study are based on the BY4741 laboratory strain (Brachmann et al., 1998) except for the PJ69-4a that were used for the Y2H assays (James et al., 1996). See the complete list of yeast strains and primers in Table S1. Cells were genetically manipulated using a transformation method that includes the usage of lithium-acetate, polyethylene glycol, and single-stranded DNA (Daniel Gietz and Woods, 2002). A pYM-based plasmid (Janke et al., 2004) was modified to contain the last 10 aa of different PTS1 proteins at the C′ of the GFP sequence. Point mutations were introduced using restriction-free cloning. Plasmids are described in Table S2. Constructs were genomically integrated into the HO locus in strains containing Pex3-mCherry, with or without *pex5* deletion and constitutive *PEX9* expression. Primers for validation of correct insertion were designed using the Primers-4-Yeast website (Yofe and Schuldiner, 2014).

### Yeast growth media

Synthetic media used in this study contains 6.7 g/L yeast nitrogen base with ammonium sulfate (Conda Pronadisa #1545) and 2% glucose, with complete amino acid mix (oMM composition, Hanscho et al., 2012), unless written otherwise. When hygromycin or geneticin antibiotics were used, media contained 0.17 g/L yeast nitrogen base without ammonium sulfate (Conda Pronadisa #1553) and 1 g/L of monosodium glutamic acid (Sigma-Aldrich #G1626) instead of yeast nitrogen base with ammonium sulfate. When mentioned, 500 mg/L hygromycin B (Formedium), 500 mg/L geneticin (G418) (Formedium), and 200mg/L nourseothricin (Silcol Scientific Equipment LTD) were used.

### Yeast library preparation using a synthetic genetic array (SGA)

To create collections of haploid strains containing GFP-tagged proteins with additional genomic modification (i.e. Pex3-mCherry (a peroxisomal marker), *PEX5* deletion *(Δpex5*), and *PEX9* constitutive-expression (*TDH3pr-PEX9*)), a query strain was constructed based on an SGA compatible strain (for further information see Table S1). Using the SGA method (Cohen and Schuldiner, 2011; Tong and Boone, 2006) the query strain was crossed with a collection of strains from the SWAT N’-GFP library (Yofe et al., 2016; Weill et al., 2018) containing ~40 strains of recently-identified peroxisomal proteins (Yifrach et al., 2021) together with controls. To perform the SGA in an arrayed format, we used a RoToR benchtop colony arrayer (Singer Instruments). In short: mating was performed on rich medium plates, and selection for diploid cells was performed on SD-URA plates containing Nourseothricin, Hygromycin, and Geneticin antibiotics. Sporulation was induced by transferring cells to nitrogen starvation media plates for 7 days. Haploid cells containing the desired mutations were selected by transferring cells to SD-URA plates containing the same antibiotics as for selecting diploid cells, alongside the toxic amino acid derivatives 50 mg/L Canavanine (Sigma-Aldrich) and 50 mg/L Thialysine (Sigma-Aldrich) to select against remaining diploids, and lacking Histidine to select for spores with an A mating type.

### Yeast library preparation from a library of pooled plasmids

A library of pooled Venus-PTS1 plasmids (DeLoache et al., 2016) was transformed genomically into a strain containing Pex3-mCherry as a peroxisomal marker, *Δpex5*, and *TDH3pr-PEX9*. 288 single colonies were picked to a 384-well plate containing SD-URA liquid media supplemented with Nourseothricin, Hygromycin, and Geneticin for selection. Then, the collected strains were imaged using automated fluorescence microscopy. In parallel, the DNA of each strain was extracted by dissolving in 20 mM NaOH followed by boiling at 95°C for 20 minutes in a PCR machine. Then, a 270 bp DNA containing the variable PTS1 region was amplified using a PCR reaction with appropriate primers (F - cgaaaagagagatcacatgg, R - gaaagcaacctgacctacag). The amplified DNA was cleaned using a GenElute 96 Well PCR Clean-up kit (Sigma-Aldrich) and sent for sequencing.

### Automated fluorescence microscopy

The collections (~40 strains of NOP1pr-GFP-recently identified peroxisomal proteins, Figure 1, and 288 strains with YFP-variable_sequence-SKL, Figure 5) were visualized using an automated microscopy setup: cells were transferred from agar plates into 384-well polystyrene plates for growth in liquid media using manual handling. Liquid cultures were grown in a LiCONiC incubator, overnight at 30°C in an SD-URA medium. An EVO freedom liquid handler (TECAN) connected to the incubator was used to dilute the strains to an OD600 of ~0.2 into plates containing SD medium (6.7 g/L yeast nitrogen base and 2% glucose) supplemented with –URA amino acids. For the ~40 NOP1pr-GFP strains we performed an additional screen in S-oleate (6.7 g/L yeast nitrogen base, 0.2% oleic acid, and 0.1% Tween-80) supplemented with –URA amino acids. Plates were incubated at 30°C for 4 hours in SD medium or for 20 hours in S-oleate. The cultures in the plates were then transferred by the liquid handler into glass-bottom 384-well microscope plates (Matrical Bioscience) coated with Concanavalin A (Sigma-Aldrich). After 20 minutes, wells were washed twice with SD-Riboflavin complete medium (for screens in glucose) or with double-distilled water (for the screen in oleate) to reduce autofluorescence, remove non-adherent cells, and obtain a cell monolayer. The plates were then transferred to the ScanR automated inverted fluorescence microscope system (Olympus) using a robotic swap arm (Peak Robotics). Images of cells in the 384-well plates were recorded in the same liquid as the washing step at 24°C using a 60× air lens (NA 0.9) and with an ORCA-flash 4.0 digital camera (Hamamatsu). Images were acquired in two channels: YFP (excitation at 488 nm, emission filter 525/50 nm) and mCherry (excitation at 561 nm, emission filter 617/73 nm). Image analysis was performed manually using ImageJ software.

### Manual microscopy

Manual microscopy imaging was performed for strains with NOP1pr-GFP-last 10 aa of Fsh3 (Figure 2) Mdh3, Cat2, and Lys1 (Figure 3). Yeast strains were grown as described above for the high-throughput microscopy with changes in the selection required for each strain (See yeast strain information in Table S1). Imaging was performed using the VisiScope Confocal Cell Explorer system, composed of a Zeiss Yokogawa spinning disk scanning unit (CSU-W1) coupled with an inverted Olympus microscope (IX83; x60 oil objective; Excitation wavelength of 488nm for GFP). Images were taken by a connected PCO-Edge sCMOS camera controlled by VisView software.

### Image analysis of peroxisomal targeting efficiency

To determine peroxisomal targeting efficiency, the GFP-PTS1 intensity was extracted from the microscope images, only for the strains that expressed Pex9 as the sole PTS1 targeting factor (Fig. 2A, right panel). The targeting efficiency was calculated as the ratio between the mean intensity in peroxisomes and the mean intensity in the cytosol. In all strains the GFP-PTS1 construct was expressed under the same constitutive promoter, *NOP1pr*, hence the total pool of GFP in the cells was expected to be similar. Since the GFP-PTS1 constructs are synthesized in the cytosol, we hypothesized that the only difference in the GFP mean intensity is a result of how efficient the peroxisomal targeting by Pex9 is.

To extract mean intensities of both peroxisomes and cytosol, image analysis was done using the ScanR software to segment the different compartments in the image. An intensity module was used on the mCherry channel (Pex3-mCherry) to segment peroxisomes. To retrieve information on the cytosol, a two-steps cell segmentation process had to be carried out. First, a cell mask was created using *Neural Network* processing (implemented in the ScanR software) on the brightfield channel. Then, an intensity module on the cell mask was used to segment the cells. Following segmentation, several parameters were extracted to a text file using ScanR, including the total and mean intensity of the GFP channel, the area and circularity of the segmented objects, and the parent object ID (the identity of the cell that corresponds to its peroxisomes).

Data analysis was performed using Python. The area and circularity of the segmented cells were used to filter out objects that do not represent single cells. The mean GFP intensity in peroxisomes was calculated by dividing the total intensity in the total area of peroxisomes. The mean intensity of the cytosol was calculated as follows:

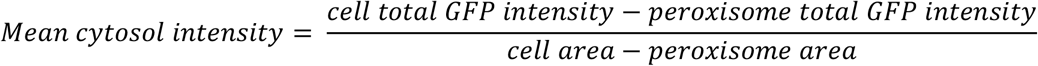

The corresponding peroxisomes for each cell were identified using the ‘parent object ID’ parameter that was extracted from the ScanR software. 207 cells were sampled from each strain. Representation of the data was done using Prism software.

### Yeast two-hybrid assay to assess protein-protein interactions

PJ69-4A cells (James et al., 1996) were transformed with one plasmid derived from pPC86 (GAL4-activation domain, AD (‘Prey’)) and pPC97 (GAL4-DNA-binding domain, BD (‘Bait’)) (Chevray and Nathans, 1992; Kerssen et al., 2006) containing genes encoding proteins of interest and selected on YNBG (0.17 % [w/v] yeast nitrogen base without amino acids, 0.5 % [w/v] ammonium sulfate, amino acids according to auxotrophic requirements, pH 6.0) plates lacking leucine (leu) and tryptophan (trp) containing 2 % [w/v] glucose (YNBG). Clones were streaked onto YNBG-trp-leu (control), YNBG -trp-leu-his-ade, and/or YNBG -trp-leu-his + 5 mM 3-amino triazole (3-AT) plates and incubated for 3, 7, or 10 days at 30 °C, respectively. HIS3 and ADE2 are under the control of GAL1 or GAL2 promoters, respectively. Thus they are only expressed when GAL-AD and GAL-BD of the bait and prey proteins are in close proximity by protein-protein interaction. 3-AT is a competitive inhibitor of the HIS3 gene product. Thus, the cells can only grow when a large amount of the HIS3 gene product is present. Here, the addition of 3-AT and longer incubation times were chosen to visualize weak protein-protein interactions by cell growth on plates lacking histidine or adenine.

### Molecular Modeling and Molecular Dynamics (MD) simulations

Model structures of Pex9 TPR (PTS1 binding) domain, residues 287-484, with bound 6 amino-acid peptides were constructed based on the experimental structures of human Pex5 complexes. The sequence identity for the TPR domain was only 29% to the human Pex5 TPR domain structure in Gatto et al., 2000 (PDB entry 1FCH) but it was spread along the whole sequence. We explored the stability of the Pex9/peptide complexes using molecular dynamics. Each starting model was immersed in a box of water, neutralized and energy minimized. Two trajectories of 100ns were calculated for each model complex, frames were extracted at 5ns intervals and inspected manually. The starting model for yeast Pex9 was constructed using Modeller (Šali and Blundell, 1993) as implemented in UCSF-Chimera (Pettersen et al., 2004). MD simulations were executed with the Gromacs package (Van Der Spoel et al., 2005). UCSF-chimera was used to visualize frames from the Pex9/peptide trajectories and to produce figure 4.

## Acknowledgments

Work in the Schuldiner lab is supported by the ERC CoG OnTarget (864068) and an Israeli Science Foundation grant 760/17. The robotic system of the Schuldiner lab was purchased through the kind support of the Blythe Brenden-Mann Foundation. MS is an Incumbent of the Dr. Gilbert Omenn and Martha Darling Professorial Chair in Molecular Genetics. RE is supported by grants of the Deutsche Forschungsgemeinschaft (FOR1905). EY is supported by the Ariane de Rothschild Women Doctoral Program. MS, RE, and LDCZ are supported by grants from the Marie Curie Initial Training Networks (from the European Commission, PerICo 812968, and PerFuMe 316723). We would like to thank Prof. John Dueber for sharing the library of pooled Venus-PTS1 plasmids with us.

## Author contribution

EY, MR, and LDCZ performed the experiments; ME performed the molecular dynamics simulations and analysis; EY and ZG performed the image analysis of Fsh3 PTS1 targeting; AT and YP generated all the plasmids that were used for the Pex9 targeting assay using microscopy; MK contributed to conceptualizing the initial work; WS, RE, MS, and EZ supervised the work; EY, MR, WS, RE, MS, and EZ wrote the manuscript; All authors read and gave feedback on the manuscript.

## Supplementary figure legends

**Figure S1.**
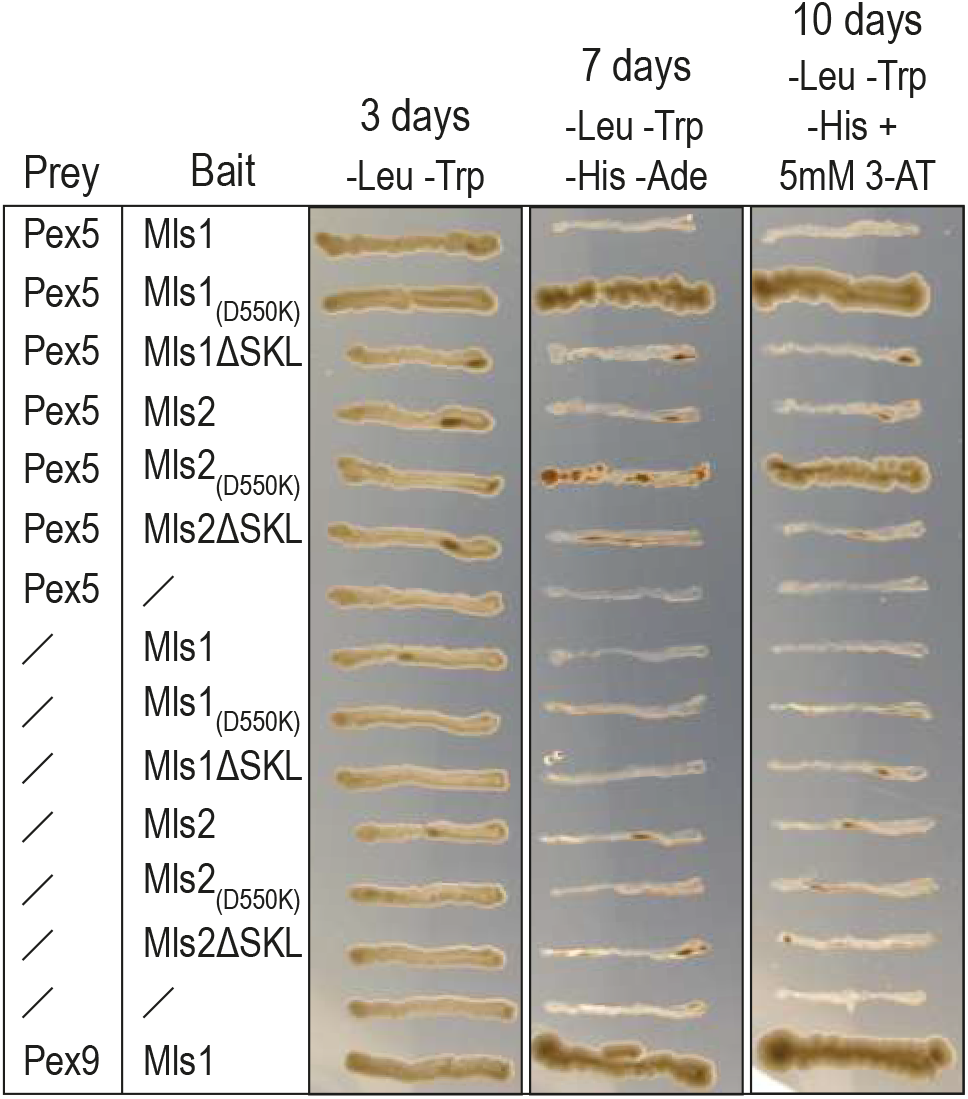
D to K substitution in Mls1 and Mls2 PTS1 contexts promote interaction with Pex5. Point mutagenesis that substitutes a negatively charged amino acid, D, with a positively charged amino acid, K, in position −5 of Mls1 and Mls2 stimulated the interaction of their PTS1 with Pex5 although the native PTS1 sequences do not interact with Pex5.

**Figure S2.**
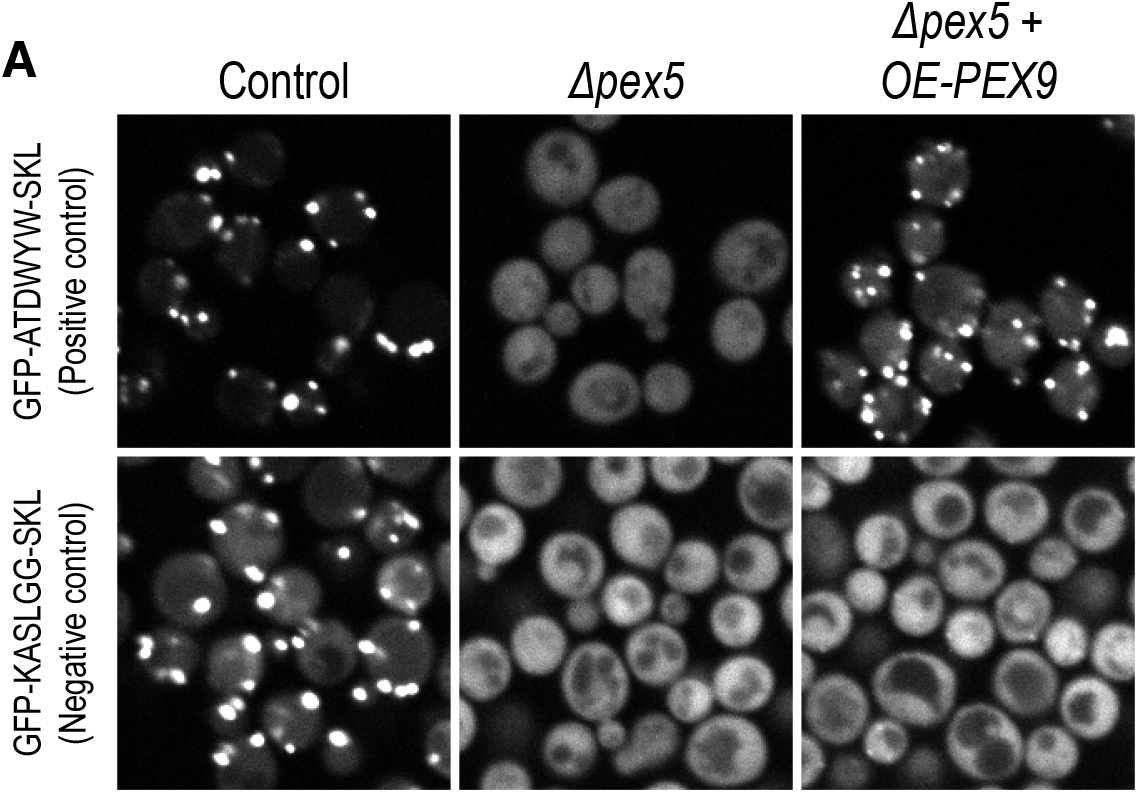
Manual validation of the YFP-XXXXXX-SKL screen results. Two sequences that showed either the highest (ATDWYW-SKL) or the lowest (KASLGG-SKL) peroxisome/cytosol localization ratio were fused to the C’ of a GFP in an integration plasmid. The manual tagging and imaging confirmed the screen results.

## Supplementary tables

**Table S1. Yeast strains and primers used in this study**.

**Table S2. Plasmids used in this study**.

